# Locus coeruleus degeneration is associated with cortical tau deposition and cognitive decline in older adults at familial risk of Alzheimer’s disease

**DOI:** 10.64898/2025.12.02.691923

**Authors:** Alfie Wearn, Kate M. Onuska, Hayley R.C. Shanks, Rahul Gaurav, Stéphane Lehéricy, Giulia Baracchini, Colleen Hughes, Jennifer Tremblay-Mercier, Sridar Narayanan, John Breitner, Judes Poirier, Taylor W. Schmitz, Gary R. Turner, R. Nathan Spreng, the PREVENT-AD Research Group

**Author notes:** Corresponding authors: Alfie Wearn or R. Nathan Spreng.

## Abstract

**INTRODUCTION:** The locus coeruleus (LC) is among the first sites of tau pathology in Alzheimer’s disease (AD) and may seed neocortical tau.

**METHODS:** We used longitudinal neuromelanin-sensitive MRI to assess LC integrity *in vivo* in a cohort of cognitively unimpaired older adults with familial risk of AD in relation to tau and amyloid PET and long-term cognitive trajectories.

**RESULTS:** We showed that both LC integrity at baseline and its rate of degeneration over time independently predicted a neocortical pattern of tau deposition. In keeping with the known function of the LC, neuropsychological tests showed that LC integrity at baseline predicted changes in attention. Finally, we found that longitudinal LC degeneration correlated with memory decline in people with elevated neocortical amyloid burden.

**DISCUSSION:** Our findings underscore the importance of LC in AD pathogenesis. Longitudinal measurement of LC degeneration may help distinguish trajectories of age-related cognitive decline and early AD.

## Background

The locus coeruleus (LC), a tiny nucleus of noradrenergic cell bodies in the brainstem’s ventral pontine area, is the among the first sites of tau neurofibrillary tangle formation in Alzheimer’s disease (AD)^1^. LC tau pathology appears decades before medial temporal lobe atrophy and onset of cognitive symptoms^2,3^. Together with other neuromodulatory subcortical nuclei of the ascending arousal system (e.g. cholinergic basal forebrain), the LC is thus a likely candidate for an origin point of tau pathology in the brain^4,5^. The long, highly branched axonal projections of these neurons may make them particularly vulnerable^5–7^, and capable of seeding tau to their cortical and subcortical targets^8,9^. LC projections are especially strong to cholinergic basal forebrain and medial temporal structures^10^, providing a plausible route for tau spread from LC to basal forebrain, entorhinal cortex and hippocampus, before more widespread neocortical involvement in the archetypical *Braak* stages^2,11^. In contrast, the LC is relatively spared from degeneration in healthy aging^12–16^. It remains debated whether reported associations between LC degeneration and age reflect true aging effects or incipient AD pathology^17–19^. Clarifying the role of LC in healthy versus pathological aging is therefore critical for understanding early AD pathogenesis and progression.

The LC noradrenergic system plays a key role in sustaining attention^20–24^. In older age, preserved LC health may underlie resilience to cognitive decline, possibly representing a substrate of cognitive reserve^25,26^. When AD pathology emerges this resilience is progressively challenged and cognitive deficits appear, typically manifesting first as episodic memory impairment^27^. Dissociating natural inter-individual variation in LC integrity from signs of pathological degeneration is thus essential for distinguishing healthy aging from AD-related pathological trajectories. One approach is to investigate how longitudinal trajectories of LC integrity relate to distinct cognitive domains (e.g., attention vs. memory).

Neuromelanin-sensitive MRI allows for sensitive and specific *in vivo* assessment of the LC health in older adults^28–30^. Neuromelanin, which is thought to sequester neurotoxic by-products of catecholamine production (e.g. noradrenaline, dopamine), is found almost exclusively in catecholaminergic cell bodies such as the LC^31,32^. Neuromelanin accrues through early and middle adulthood, peaking around age 60^33–36^. Thereafter, LC neuromelanin signal correlates closely with neuronal density in both unimpaired older adults and people with AD^37,38^, and can therefore be considered a marker of structural integrity. Neuromelanin MRI-based measures of LC integrity appear to predict neocortical tau pathology^39–42,37^ and may be useful for tracking disease onset and progression.

To date, almost all studies of LC integrity have been cross-sectional, and none have assessed whether the longitudinal rate of LC degeneration provides superior or additional information about pathology over and above a single measure of integrity. Here, we leveraged state-of-the-art neuroimaging, including neuromelanin MRI and amyloid and tau PET, together with longitudinal multidomain neuropsychological testing to study how changes in LC health may relate to other hallmarks of AD pathology. We studied a cohort of older adults with a strong family history of AD, but who were nonetheless cognitively unimpaired at baseline. We tested the following hypotheses: 1) LC integrity would be lower in older (vs. younger) adults across the entire group, but this association would be explained by the presence of incipient AD pathology; 2) lower baseline LC integrity and steeper rates of LC degeneration would be associated with greater neocortical tau deposition in regions corresponding to early Braak stages, especially in individuals with higher amyloid burden; and 3) lower baseline LC integrity and steeper rates of LC degeneration would be associated with steeper trajectories of cognitive decline, showing associations in the specific domains of memory and attention.

## Methods

### Participants

We used data from PREVENT-AD, a longitudinal cohort study of older adults with a parent or multiple siblings with clinically diagnosed AD^43,44^. After exclusions detailed below, we studied 199 participants (mean age: 68.3 ± 5.2, 69.8% female) having multiple available timepoints of neuromelanin MRI and valid cognitive test data. Participants were followed up for an average of 46.5 ± 14.9 months. A subsample of 178 participants (mean age: 68.3 ± 5.1, 70.0% female) also had PET assessment of amyloid and tau on at least one timepoint, as described below.

All participants gave informed written consent, and all PREVENT-AD study features were approved by the McGill institutional review board and/or the *Comité d’éthique de la recherche du CIUSSS de l’ouest de l’ile de Montréal*. All procedures complied with the ethical standards of the 1964 Declaration of Helsinki.

### MRI Acquisition

MRI scans were acquired on a 3T Siemens PrismaFit at the Douglas Research Centre, including a T1-weighted anatomical scan (Magnetization-Prepared Rapid Acquisition Gradient Echo (MPRAGE), 1 mm^3^ isotropic resolution, TR/TE/TI = 2300/2.96/900 ms, FA = 9°, TA = 5:30), and a neuromelanin-sensitive axial slab of the brainstem and midbrain: 0.7x0.7x1.8mm, TR/TE=600/10 ms, FA=120°, TA=8:27.

MRI was collected in three distinct waves between 2019-2024. Participants (n=269) had between one and three scans including neuromelanin MRI. Of these, 207 participants had at least two timepoints of neuromelanin MRI data and were therefore included in this study (mean interscan interval: 28.8 ± 6.4 months; mean total follow-up time: 46.5 ± 14.9 months; Average number of scans: 2.6 ± 0.49.

### PET Acquisition

Participants underwent amyloid ([^18^F]NAV4694), and tau ([^18^F]flortaucipir) PET. Scans were acquired on a brain-dedicated high-resolution research tomograph (HRRT) scanner at the McConnell Brain Imaging Centre of the Montreal Neurological Institute-Hospital.

For amyloid PET, 6 mCi radiotracer was injected and the scan was acquired in the time window 40-70 minutes post-injection. For tau PET, approximately 10 mCi was injected and the scan was acquired in the time window 80-100 minutes post-injection. Frames of 5 minutes were acquired. Images were reconstructed using an iterative reconstruction method (OP-OSEM, 10 iterations, 16 subsets). Images were motion and decay corrected, and scatter correction was performed using a 3D method.

Two waves of PET data collection occurred, the first in 2016-2017, the second between 2019-2023. For participants with multiple PET scans, images were processed by aligning the temporally closest PET-MRI scans to minimize anatomical differences between images; however, in order to maximize our sensitivity to AD pathology, we related demographic, behavioral and MRI variables to PET measures from participants’ latest PET session, which was often in-line with the second wave of MRI collection. Therefore, for all analyses baseline MRI preceded PET collection by an average of 208 ± 528 days.

### Image Processing

#### Measuring LC Integrity – Neuromelanin MRI

At each timepoint, we calculated the ratio of neuromelanin-MRI signal intensity in the LC to that of an adjacent pontine reference region as a marker of structural integrity (shown to correlate with neuronal density^37,38^). We calculated LC integrity using an automated pipeline described previously^45^. Briefly, *a priori* defined standard space masks of the LC and immediate surrounding area were transformed and resliced into a given timepoint’s neuromelanin scan’s space. These masks (separate for left and right hemispheres) were used as search spaces in which to identify the 10 brightest contiguous voxels, which corresponded to LC (confirmed by visually checking all automatically produced masks). Scans with mislabeled regions or artefacts on the region of interest were excluded from further analysis. A total of 24 sessions across 21 participants were excluded due to masking errors or image artefacts. Following the exclusion of these timepoints, eight participants no longer had two timepoints of valid LC measurement and were therefore excluded from the main sample (total sample size after exclusions: 199). A pontine tegmentum reference region mask was also transformed to native space in the same way. Relative intensity of the LC (LC_RI_) was calculated by dividing the LC signal intensity by the reference region signal intensity, multiplying by 100 and averaging over all slices. Left and right LC intensities were averaged for all analyses.

#### Measuring AD pathology – PET

PET processing was performed in MATLAB (v2024b) using SPM (v25.01.02) and the CAT12 (v12.9) toolbox, using this pipeline: https://github.com/hayleyshanks/Longitudinal-MRI-PET-preproc ^46^). We previously created an MRI population template representative of older adult brains using 3T T1-weighted data from 424 participants from the Alzheimer’s Disease Neuroimaging Initiative (ADNI) cohort as described in^46^. First, each MPRAGE image was linearly co-registered to the ADNI template brain using SPM’s Estimate tool. These images were then segmented into grey matter (GM), white matter (WM) and CSF compartments using CAT12. Mean image quality rating was 86.6% ± 0.20%. No segmentations fell below a threshold of 70% that warranted further examination. GM and WM segmentation images (rigid-aligned to template) were then used to calculate non-linear warps to the ADNI population template using geodesic shooting^47^.

All frames of PET images (four for tau, six for amyloid) were aligned to the final image of the series to account for between-volume motion and then averaged. The PET image was then co-registered using SPM’s Estimate & Reslice tool to its temporally closest MPRAGE (native-space). PET data was then warped to the population template brain using warps previously calculated using the MPRAGE images. Following this process, two participants were excluded for having poor alignment with the MPRAGE. We then calculated standardized uptake value ratio (SUVR) for all brain voxels relative to inferior cerebellum gray matter for tau and whole cerebellum gray matter for amyloid. Partial volume error correction was performed for both tracers and was calculated using Müller-Gartner method with the PETPVE toolbox^48^ (default settings except GM threshold was changed from 0.5 to the slightly more liberal 0.3).

To assess global amyloid burden in each individual, we calculated a volume-weighted mean of SUVR in the following brain cortical regions from the Desikan-Killany atlas: Caudal Middle Frontal, Lateral Orbitofrontal, Medial Orbitofrontal, Pars Opercularis, Pars Orbitalis, Pars Triangularis, Rostral Middle Frontal, Superior Frontal, Frontal Pole, Inferior Parietal, Precuneus, Superior Parietal, Supramarginal, Caudal Anterior Cingulate, Posterior Cingulate, Isthmus Cingulate, Rostral Anterior Cingulate, Middle Temporal, Superior Temporal, Inferior Temporal.

The volume-weighted mean was calculated according to the following formula:

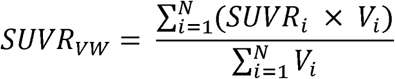

Where *SUVR_i_* is the *SUVR* of region *i*, V*_i_* is the corresponding volume of region *i*, and *N* is the number of regions included.

### Assessing Cognition

Cognitive performance was assessed using the Repeatable Battery for the Assessment of Psychological Status (RBANS)^49^. This 30-minute battery, which is designed for characterizing cognitive trajectory in older adults, comprises tests of five cognitive domains: attention, memory (immediate), memory (delayed), language, and visuospatial/constructional ability. These domain scores can be used individually or combined into an omnibus score of general cognitive ability. Raw, non-age-adjusted scores were used here.

To more accurately characterize individuals experiencing cognitive decline we leveraged the full range of PREVENT-AD RBANS data collected between 2012 and 2024 (by contrast, neuromelanin MRI data collection began in 2019). We therefore assessed long-term trajectories of largely retrospective cognitive performance relative to the MRI baseline timepoint. On average, each participant had 7.54 ± 2.33 RBANS assessments (range: 2-11) over a total of 7.96 ± 1.67 years (range: 5-10). Cognitive data were typically collected within one day of MRI scanning.

### Statistical Analysis

#### Assessing within- and between-subject correlates of age and longitudinal change in LC

The linear relationship between LC integrity at baseline and its longitudinal rate of decline was assessed using liner regression using the ‘*lm*’ function in R v4.4.2, correcting for age, sex and education.

To test whether longitudinal change in LC was significant within individuals and whether LC integrity at baseline or its change varied by age, we ran a longitudinal mixed effects model (using ‘*lmerTest*’ v 3.1-3^50^ in R) defined as:

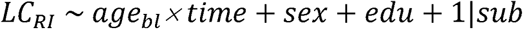

where *age_bl_* denotes age at baseline, *edu* denotes years of education and *1|sub* represents a random intercept for each subject. Main effects on interaction terms are implicitly defined. A significant effect of age at baseline would indicate an overall between-subject effect of age. A significant effect of time would indicate within-subject longitudinal change. Finally, a significant age × time interaction would indicate that within-individual change was different across different ages.

To test whether the age effect was an artefact of incipient AD, we repeated the analysis adding global amyloid burden as an extra interaction term as follows:

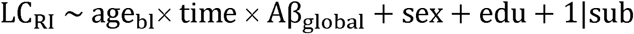

#### Testing the association between LC integrity and cortical tau deposition

We aimed to test whether LC integrity at baseline (bl-LC_RI_) or rate of degeneration (slope of longitudinal change in LC_RI_, defined as ΔLC_RI_) was related to neocortical tau deposition, and whether the association was stronger in people with greater amyloid load. ΔLC_RI_ was defined as the β-coefficient of the regression slope between time and LC_RI_ for each participant. Using SPM12’s Multiple Regression framework^51^, we defined the following model:

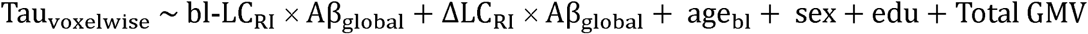

where *Total GMV* is gray matter volume, calculated from the CAT12 ‘Segment’ step^52^.

The analysis was run across all voxels of the template-space tau SUVR brain image. Significant clusters of voxels were identified in voxel space using threshold free cluster enhancement (TFCE, r269) (5000 permutations, E=0.5, H=2.0)^53^. Significant voxels (family-wise error corrected p<0.05 for robust correction for multiple comparisons) were then projected to surface for presentation.

#### Testing the association between LC integrity and cognition

Finally, we tested whether long-term trajectories of cognition were predicted by bl-LC_RI_ or ΔLC_RI_ and whether any such association was modulated by amyloid burden using the following linear mixed-effects models:

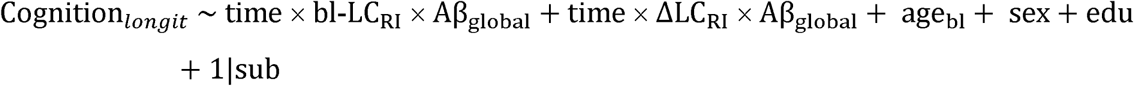

where the dependent variable is longitudinal cognition for either RBANS total score, or scores of one of the five cognitive subdomains.

## Results

### Longitudinal changes in LC

Across the cohort, bl-LC_RI_ values averaged 119.7 ± 3.1% and changed by −0.187 ± 0.630% per year. We observed a negative association between bl-LC_RI_ and longitudinal change in LC (ΔLC_RI_) after correcting for age, sex and education (n=199, t=-2.71, p=0.007, Figure 1C). Our mixed-effects analysis revealed that LC_RI_ declined significantly over time within individuals (t_336_ = −4.87, p<0.0001), and was lower in older individuals (t_262_=2.01, p=0.045) (Figure 1D). We did not observe a significant age × time interaction (Figure 1E), indicating that rate of LC degeneration did not change as function of age (t_342_=-1.04, p=0.301). No significant differences were observed with sex (t_256_=0.589, p=0.556) nor with years of education (t_257_=1.33, p=0.186).

**Figure 1.**
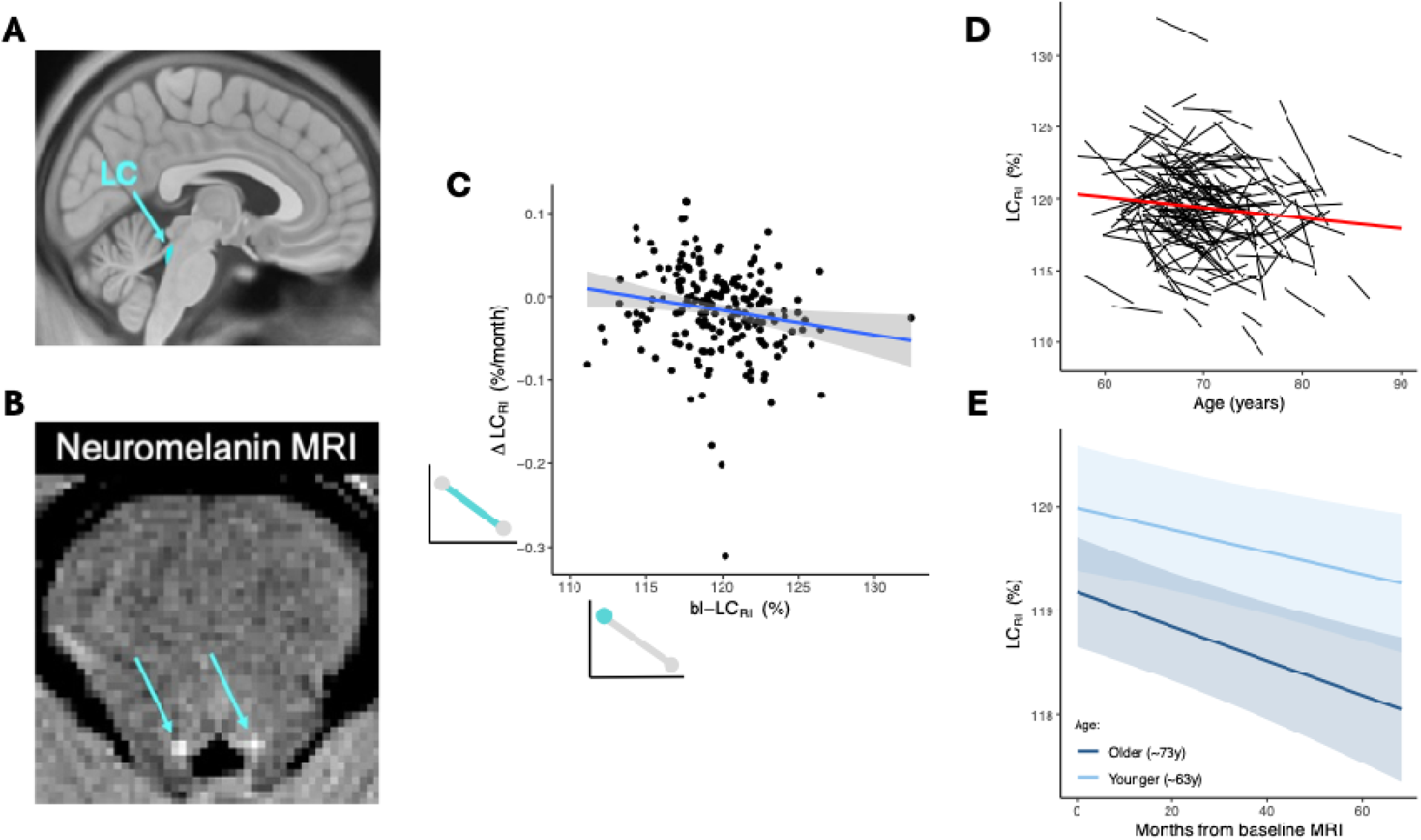
LC visualisations, longitudinal change and association with age. A) LC anatomical placement on sagittal MNI-space brain. LC mask shown is from Ye atlas^54^ for visualisation only. B) Neuromelanin MRI in axial plane of a single participant with LC hyperintensities indicated by arrows. C) Association between LC integrity at baseline and its rate of degeneration. The panel includes small schematics representing LC baseline and longitudinal change adjacent to their respective axes of the main plot. These schematics are used in figures throughout this paper. D) Longitudinal trajectories of LC integrity as a function of age in individual participants. Group trend line shown in red. E) Longitudinal trajectories of LC integrity in younger (∼63y) vs older (∼73y) individuals. In the model, the interaction term for age was non-significant, which is apparent in the figure as slopes are similar between age groups. Note that age is displayed here using binary groups for visualisation purposes only; analyses incorporated age as a continuous variable in all instances (‘younger’ group here is represented as 1 SD below the mean; ‘older’ as 1 SD above the mean).

We next added global amyloid burden as an interaction term to test whether the age effect remained apparent independent of amyloid load. Here we still observed a significant within-subject longitudinal change (t_307_=-4.27, p<0.0001), but the between-subject age effect was no longer apparent (t_236_=-1.12, 0.265), indicating that those with incipient AD were driving the groupwise above-noted relationship between age and LC_RI_ in the previous analysis. There was no significant main effect of global amyloid (t_386_=-0.928, p=0.354) on LC_RI_, nor any interaction between amyloid and age (age × amyloid: t_369_=-0.721, p=0.472). Further, we found no interaction between age, amyloid and rate of LC degeneration (age × time × amyloid: t_325_=0.630, p=0.529).

### LC integrity at baseline and rate of degeneration independently predict neocortical tau deposition

We observed strong associations between global amyloid burden and voxelwise neocortical tau deposition with regional specificity in temporal (including parahippocampal gyrus), parietal and lateral occipital areas (n=178; Figure 2A). Main effects for bl-LC_RI_ and ΔLC_RI_ did not reveal any regions of significant association with tau burden across the cortex. We did, however, observe significant interactions of these terms with amyloid burden (Figure 2B & C). We interpret this observation as suggesting that in people with greater amyloid burden, both lower bl-LC_RI_ (Figure 2B) and more negative ΔLC_RI_ (Figure 2C) predict neocortical tau deposition.

**Figure 2.**
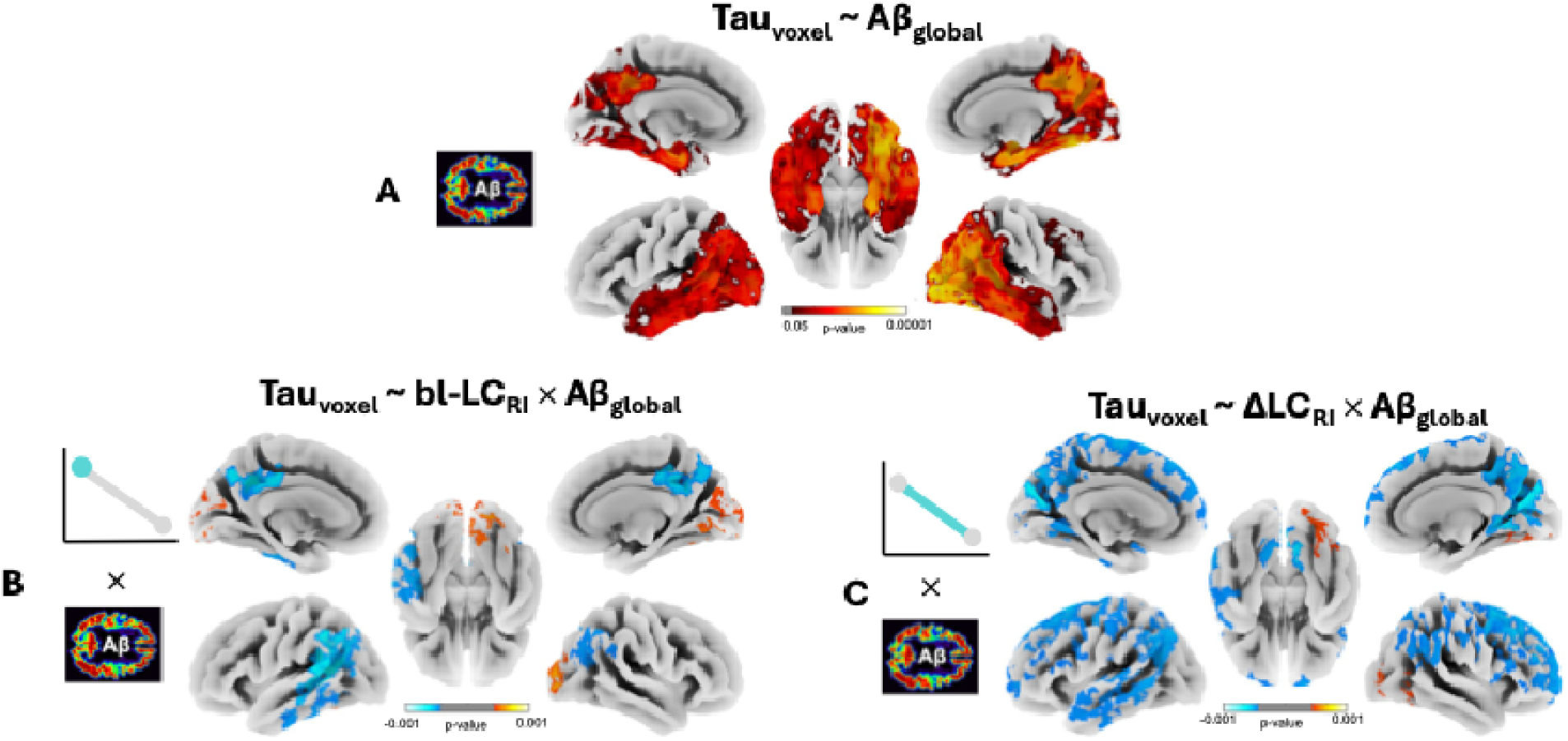
Surface-projected statistical maps of main effects predicting neocortical tau. A) Regions where global amyloid (Aβ_global_) predicted local neocortical tau burden. In regions shown in red-yellow, elevated Aβ_global_ predicted greater neocortical tau. B) Regions where the bl-LC_RI_ × Aβ_global_ interaction was statistically significant. In blue regions, lower bl-LC_RI_ predicted greater tau deposition in people with elevated Aβ_global_. In red-yellow regions (occipital cortices) greater bl-LC_RI_ predicted more tau deposition in people with elevated Aβ_global_. C) Regions where the ΔLC_RI_ × Aβ_global_ interaction is statistically significant. In blue regions, steeper decline in LC_RI_ predicted greater tau deposition in people with elevated Aβ_global_. In red-yellow regions (occipital cortices) steeper decline in LC_RI_ predicted less tau deposition in people with elevated Aβ_global_. Each main effect is again denoted by a schematic to the left of the respective panel. Throughout the plots, positive associations are denoted in warm colours, and negative associations in cool colours. ‘Negative’ p-values on colour scales indicate p-values of the negative association. All panels show FWE-corrected TFCE log p-value maps where p<0.05. Unthresholded maps are shown in Fig. S1 in Supplementary Material. Main effects of bl-LC_RI_ and ΔLC_RI_ are not shown as there were no statistically significant voxels.

Among those with higher global amyloid, low bl-LC_RI_ predicted greater tau deposition in lateral temporal and medial parietal cortices, but predicted lower tau deposition in occipital regions. Again in people with higher global amyloid, more negative slopes of ΔLC_RI_ predicted greater tau deposition broadly across lateral temporal cortices (in left hemisphere only), medial and lateral parietal cortices, and across frontal regions. Some sparse areas of the inverse association were also observed in occipital cortices, such that more negative slopes of ΔLC_RI_ predicted lower tau deposition.

These results were robust to various choices in statistical analytic approach and processing including: transforming voxelwise tau across individuals to reduce skew, not performing PVC of amyloid or tau, and including global amyloid as a binary positivity variable, at a 30-centiloid threshold^55,56^ (Supplementary Material, Figs. S2-4).

### LC integrity at baseline predicts global but not voxelwise amyloid deposition

Neither bl-LC_RI_ nor ΔLC_RI_ predicted voxelwise amyloid SUVR across any region of the cortex (n=178). However, a linear regression analysis did reveal a significant association between global amyloid and bl-LC_RI_ (t=-2.31, p=0.021) but not ΔLC_RI_ (t=0.328, p=0.743).

### LC integrity at baseline and rate of degeneration predict cognitive decline

We studied baseline and longitudinal associations between LC integrity and cognitive decline, testing also whether any such associations were stronger in people with elevated global amyloid (n=178; Figure 3). The RBANS total composite score declined significantly over time in the whole cohort (main effect of time: t_1201_=-6.92, p<0.0001) but were significantly lower in people with elevated global amyloid at baseline MRI (main effect of amyloid: t_186_=-2.94, p=0.004). Cognitive decline was also accelerated in persons with higher global amyloid (time × amyloid interaction: t_1204_=-5.35, p<0.0001).

**Figure 3.**
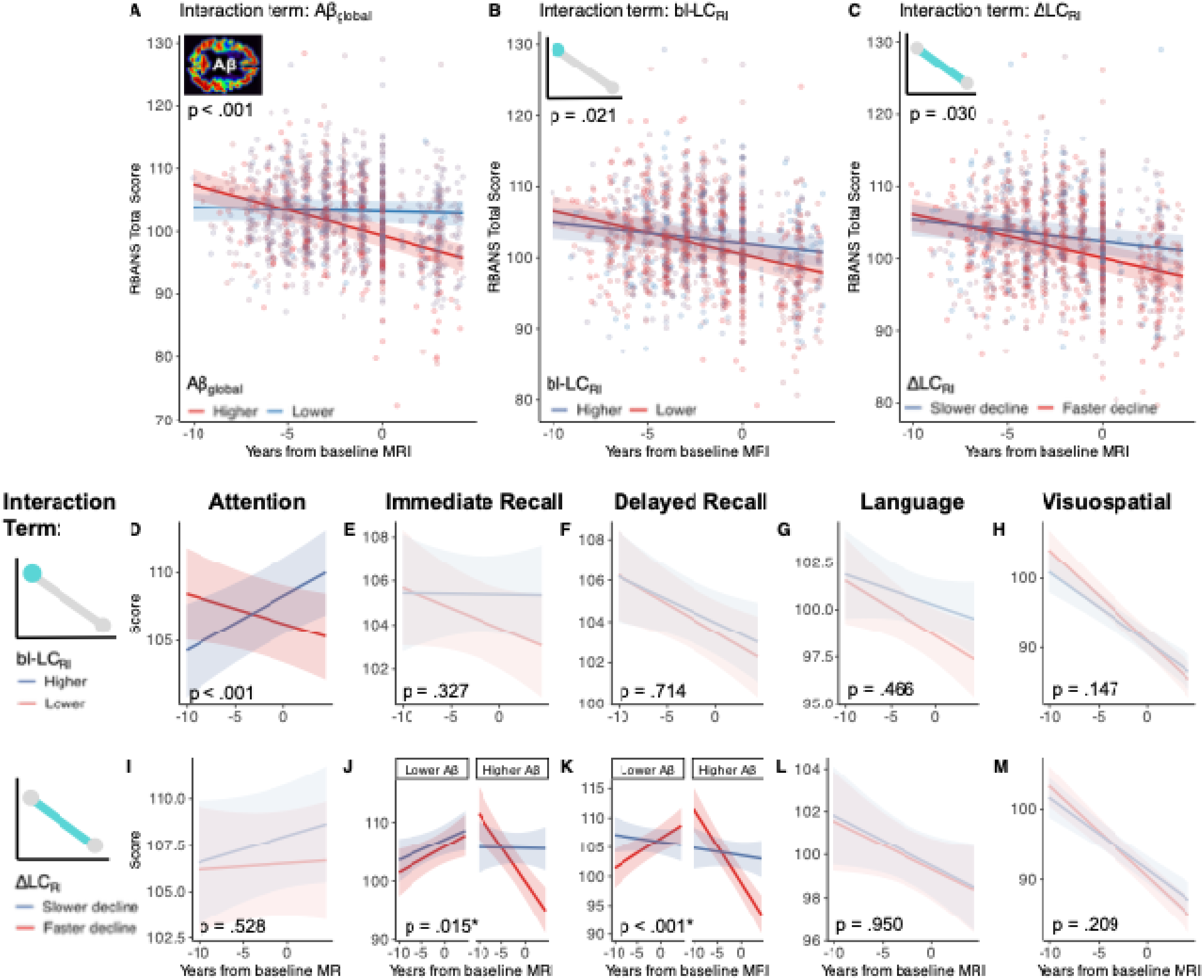
Longitudinal trajectories of cognitive decline modulated by global amyloid and LC integrity. A-C) Scatterplots showing steeper cognitive decline, measured by RBANS total score in people with A) elevated global amyloid (Aβ_global_), B) lower LC integrity at baseline (bl-LC_RI_), and C) faster LC degeneration (more negative ΔLC_RI_). Points in A-C show partial residuals from a single linear mixed effects model correcting for, age at MRI baseline, sex and education. D-H) Trendlines showing associations between bl-LC_RI_ and individual cognitive domain scores. Lower bl-LC_RI_ was associated with steeper decline of the attention score only. I-M) Trendlines showing associations between ΔLC_RI_ and individual cognitive domain scores. Lower ΔLC_RI_ was associated with steeper decline of both immediate and delayed memory scores in people with elevated Aβ_global_. Small inset schematics are shown to visually denote the main effects of interest (either Aβ_global_, bl-LC_RI_ or ΔLC_RI_) in respective plot groups. P-values for the interaction between the relevant variable of interest and time are shown on each plot, with J & K showing p-values for the 3-way interaction with amyloid (denoted by *). Note that all variables were modelled continuously, but are plotted here as binary high/low trendlines (± standard error) at one standard deviation above or below the mean for visualisation purposes only. Colours are numerically reversed in A compared to other plots to align with the notion of ‘red=worse’, ‘blue=better’. Colors are faded where the main interaction effect of interest is not statistically significant. Plots are shown faceted by Aβ_global_ if the three-way interaction between LC × time × amyloid was statistically significant. LC = locus coeruleus, RI=relative intensity.

We did not observe any association between overall cognitive performance (RBANS total score) at baseline MRI with bl-LC_RI_ (t_184_=1.14, p=0.255) or ΔLC_RI_ (t_190_=1.69, p=0.094). We did, however, note steeper trajectories of cognitive decline in people with lower bl-LC_RI_ (time × bl-LC_RI_ interaction: t_1217_=2.31, p=0.021) and more negative ΔLC_RI_ (time × ΔLC_RI_ interaction: t_1216_=2.18, p=0.030). The association between bl-LC_RI_ and cognitive decline was not stronger for people with elevated global amyloid (time × bl-LC_RI_ × amyloid interaction: t_1224_=1.03, p=0.304). The association between ΔLC_RI_ and cognitive decline appeared slightly stronger among people with higher global amyloid, but this did not reach statistical significance (time × ΔLC_RI_ × amyloid interaction: t_1236_=1.87, p=0.06).

We next tested whether either bl-LC_RI_ or ΔLC_RI_ predicted longitudinal change in RBANS subscales. Lower bl-LC_RI_ predicted decline in attention performance (time × bl-LC_RI_: t_1199_=3.544, p=0.0004). More negative ΔLC_RI_ (faster LC degeneration) predicted more negative trajectories of immediate memory (time × ΔLC_RI_: t_1253_=2.86, p=0.004), an effect which was stronger in persons with higher global amyloid compared to those with lower global amyloid (time × ΔLC_RI_ × amyloid: t_1280_=2.42, p=0.016). More negative ΔLC_RI_ slopes also predicted faster declines in delayed memory in people with elevated global amyloid (time × ΔLC_RI_ × amyloid: t_1272_=4.307, p<0.0001). Language and visuospatial scores were not related to either bl-LC_RI_ or ΔLC_RI_. Full statistics for these models are shown in Supplementary Material (Table S1-2).

## Discussion

In a cohort of older adults with a familial history of AD, we examined longitudinal changes in LC integrity measured *in vivo* using neuromelanin-sensitive MRI. We observed lower LC integrity in older individuals but found that this was likely driven by incipient AD pathology. Among participants with higher global amyloid, both lower LC integrity and a faster rate of LC degeneration were independently associated with greater neocortical tau deposition in regions corresponding to Braak stages IV/V. At the same time, we observed that lower LC integrity and faster degeneration were associated with less tau deposition in occipital regions, suggesting a specificity of tau deposition pattern to LC degeneration. Finally, we showed independent associations between LC integrity and rate of LC degeneration with long-term trajectories of cognitive decline such that LC baseline integrity was associated with changes in attention, whereas rate of LC degeneration was associated with memory decline.

### LC age-related degeneration explained by incipient AD pathology

We observed a robust within-individual decline in LC integrity over an approximately 4-year follow-up period. This was accompanied by a group-wide negative association with age, whereby the older individuals (∼80y) had lower LC integrity than younger participants (∼60y). In keeping with a previous study^40^, we did not observe an interaction between age and time, which would have been expected if rate of LC degeneration increased with advancing age. In contrast, another longitudinal study of a similar size reported a significant association between rate of LC degeneration and age^57^. This discrepancy may reflect our explicit control for baseline LC integrity. The association between LC integrity and LC degeneration was weak (equivalent to r∼-0.2 according to our data) but should nonetheless be noted.

Age-related LC decline has been reported previously. Neuronal counts in the LC across the lifespan has been estimated to decrease by 20-40% (see review^20^). Neuromelanin-sensitive MRI studies report an inverted-U shaped lifespan trajectory, with LC signal peaking around age 60^33–36^. However, these exclusively cross-sectional studies do not test explicitly whether there is a robust decline in older age or whether the trajectory is better described as a plateau. Indeed many studies focusing on older adults report no age effect^12,15,16,58^. Where age effects are reported, it is often unclear whether LC changes reflect ‘healthy aging’ or incipient AD. Such studies of LC changes in ostensibly healthy aging rarely exclude individuals with AD pathology elsewhere in the brain (see ^59^ for one exception). This risks inclusion of participants with pre-symptomatic AD as an explanation of LC degeneration in some cases. In our analyses, adjusting for global amyloid burden removed the group-wise association with age while within-individual declining trajectories remained significant. This pattern supports a model in which LC integrity does not decline in healthy aging and in which marked atrophy is instead a specific feature of early AD pathology. This observation justifies our focus on disentangling the independent effects of baseline LC integrity and subsequent degeneration on downstream pathology.

### LC integrity at baseline and rate of LC degeneration independently predict neocortical tau

We found that both lower LC integrity and faster rate of LC degeneration were independently associated with greater neocortical tau deposition in people with higher global amyloid burden. In persons with elevated amyloid, associations between cross-sectional LC integrity and neocortical tau were localized to medial parietal cortices and left lateral temporal cortices, while associations between LC rate of degeneration and neocortical tau exhibited a more widespread pattern including frontal cortices. These regions roughly correspond to Braak stages IV/V^2,11^. Notably, we did not observe a significant association localized to entorhinal or parahippocampal cortices after accounting for the predictive power of global amyloid in these regions. The pattern is similar to a previous study in which LC integrity predicted tau deposition primarily in posterior cortices in people with autosomal-dominant AD (symptomatic and non-symptomatic)^41^. Our observed pattern is more extensive than some previous reports of exclusively medial-temporal localized associations between LC neuromelanin signal and tau SUVR^39,40^. Another study measured tau directly in LC, showing associations with neocortical tau in people with more global amyloid accumulation, with effects localized to Braak stage III/IV regions (limbic cortices)^60^. Thus, although there is some variability across findings, the presence of an association between LC measures and neocortical tau in individuals with likely incipient AD (by global neocortical amyloid) appears robust.

Our data are consistent with the increasingly supported model that the LC facilitates the spread of tau to the neocortex, thereby driving AD progression. In this model, the LC seeds tau initially to other neuromodulatory nuclei including the cholinergic basal forebrain, followed by the medial temporal lobes^4,5,61–64^. Later, when interacting with amyloid-induced neuronal hyperexcitability, tau is spread more rapidly and broadly across the neocortex^65–70^. According to our data and prior work^39,41,60^, existing neuroimaging tools reliably capture LC degeneration *in vivo*, and this degeneration occurs prior to clinically detectible cognitive decline but after substantial neocortical tau spread. Moreover, within-individual rates of LC degeneration seem particularly indicative of widespread tau pathology across regions in later *Braak* stages. Our finding therefore supports *in vivo* longitudinal measurement of LC integrity as a marker of tau spread and overall pathological extent. Direct measurement of tau with PET provides greater specificity but is invasive, substantially more expensive and less repeatable.

This model may, however, primarily describe the canonical pattern of tau deposition in AD. Increasing evidence indicates that AD is unlikely to follow a single trajectory, but instead comprises distinct subtypes^71–74^. These subtypes can be characterized by variation in tau distribution patterns (e.g. temporal- vs posterior-predominant) as well as in clinical presentation (e.g. memory- vs visual-predominant). We report an unexpected association whereby greater LC degeneration was related to less tau in occipital cortices. Our findings lead us to hypothesize that a posterior-predominant tau deposition subtype may have a partially non-LC-related etiology. The reverse pattern was not observed in a previous analysis of LC integrity as a predictor of neocortical tau deposition in people with autosomal-dominant AD^41^, a condition more likely to follow a unified pathological trajectory than sporadic AD. In any case, further investigation and explicit testing of this hypothesis are warranted.

### LC integrity at baseline and rate of LC degeneration independently predict changes in cognition

We found that individuals with lower LC integrity at baseline and those with a faster rate of LC degeneration exhibited steeper trajectories of cognitive decline over an approximately 14-year period, assessed mostly retrospectively. This effect was independent of global amyloid burden. As with our findings on neocortical tau distribution, this analysis demonstrates distinct and additive associations of both LC integrity and rate of LC degeneration with cognitive decline. These data are consistent with previous studies showing associations between LC health and cognition^39,57,58^, and support the notion that LC neuronal density may confer cognitive reserve and is therefore critical for the maintenance of cognitive function in older age (for a review on this topic see ^20^).

We observed that baseline LC integrity predicted the rate of change in attention over time, in line with an understanding that the primary function of LC is to sustain and modulate attention^20–24^. Our finding supports a previous study showing that LC degeneration is associated with reduced practice effects^75^ and is consistent with a model whereby LC integrity confers cognitive reserve^25,26^. Variation in LC integrity appears to correspond with the ability to maintain effective attention in cognitively unimpaired older adults^20,22,23,25,26^. We also found that the rate of LC degeneration predicted the rate of memory decline (both immediate and delayed recall), and that this effect was strongest in people with higher amyloid burden, in keeping with our hypothesis. This supports the notion that those exhibiting significant LC degeneration are most likely to be in preclinical stages of AD, which are characterized in early stages by selective impairments in memory^27,76^. One previous study reported a similar effect whereby LC integrity moderated the association between memory change and amyloid but did not examine specific tests of attention^39^. To our knowledge, ours is the first study to report specific associations of LC integrity and its rate of change with separate cognitive subdomains. Overall, our findings support the use of LC integrity and degeneration for differentiating individuals in the preclinical stages of AD from those on a healthy aging trajectory.

### Conclusions

The LC may be the earliest system directly involved in AD pathogenesis. With the development of tools that can probe LC health and neocortical tau and amyloid *in vivo*, we are now able to characterize and track earlier stages of AD than what was possible using purely histological methods. We hope that our study encourages additional research in longitudinal *in vivo* measurement of the health of this important neuromodulatory system. Future work should assess whether improving LC health can produce downstream effects on disease outcomes^77^.

Furthermore, specificity of LC degeneration can be explored in relation to atypical AD subtypes and other neurodegenerative diseases. Finally, a better understanding of the cholinergic, serotonergic, orexinergic and dopaminergic systems, including how they interact, spread tau and support compensatory mechanisms in the brain, is essential to fully understand AD pathogenesis. This line of research may ultimately support the development of a new generation of highly effective disease-modifying treatments.

## ACKNOWLEDGMENTS

We would like to thank the PREVENT-AD research team and study participants for their time and dedication in collecting the data used in this study. A full list of PREVENT-AD authors and collaborators can be found <u>here</u> and further information about the program can be found <u>here</u>. We would also like to thank the Laboratory of Brain and Cognition at the Montreal Neurological Institute for productive discussions and helpful feedback.

## CONSENT STATEMENT

All human subjects in this study provided informed written consent prior to participation.

