## Supplemental Figures for "Locus coeruleus degeneration is associated with cortical tau deposition and cognitive decline in older adults at familial risk of Alzheimer’s disease"

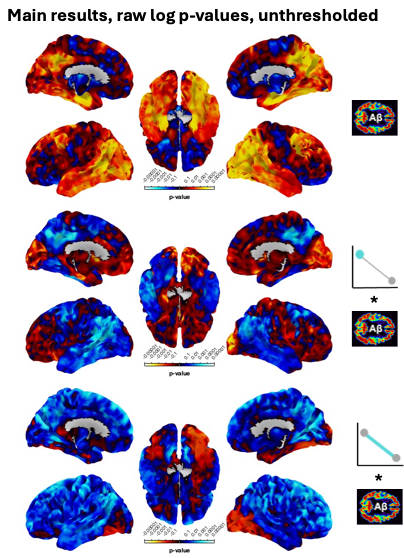

Fig. S 1 | Raw, unthresholded p-values of the effects shown in the main body of manuscript (Fig. 2). Top row: Main effect of global amyloid; Middle row: Main effect of bl-LC * global amyloid interaction term; Bottom row: Main effect of ΔLC * global amyloid interaction term.

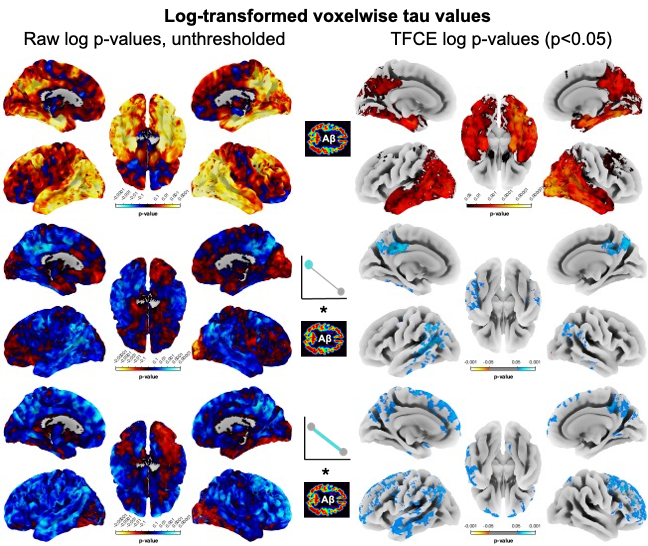

Fig. S 2 | Raw, unthresholded p-values and significant TFCE, FWE-corrected clusters of the same analysis as presented in Fig. 2 of the main paper, with the exception that voxelwise tau value were first log-transformed in order to reduce the effect of outlying values on the observed effect. Top row: Main effect of global amyloid; Middle row: Main effect of bl-LC * global amyloid interaction term; Bottom row: Main effect of ΔLC * global amyloid interaction term.

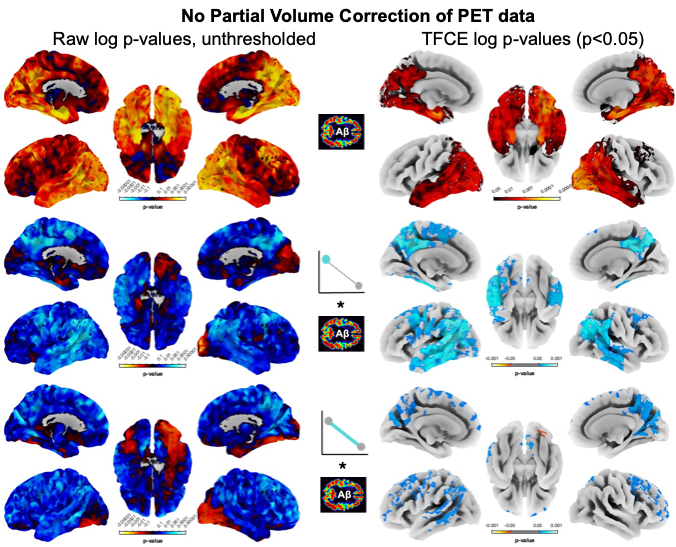

Fig. S 3 | Raw, unthresholded p-values and significant TFCE, FWE-corrected clusters of the same analysis as presented in Fig. 2 of the main paper, with the exception that amyloid and tau SUVR values were not corrected for partial volume correction (PVC). This step is often recommended but sometimes omitted as can introduce bias. For completion, we explored not conducted PVC, but see a largely similar cluster pattern. Top row: Main effect of global amyloid; Middle row: Main effect of bl-LC * global amyloid interaction term; Bottom row: Main effect of ΔLC * global amyloid interaction term.

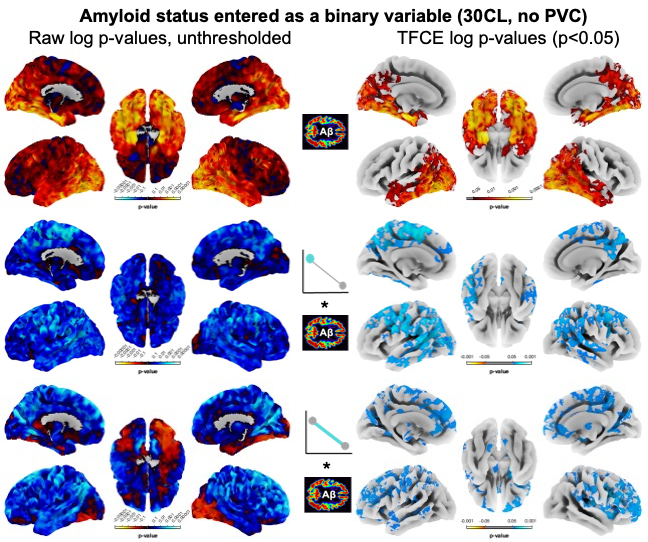

Fig. S 4 | Raw, unthresholded p-values and significant TFCE, FWE-corrected clusters of the same analysis as presented in Fig. 2 of the main paper, with the exception that partial volume correction was not applied and amyloid variables were entered as a binary measure, thresholded at 30 centiloids. This threshold is recommended as an upper limit for certainty of abnormal amyloid levels (Collij et al. 2024, Alzheimers Dement). Top row: Main effect of global amyloid; Middle row: Main effect of bl-LC * global amyloid interaction term; Bottom row: Main effect of ΔLC * global amyloid interaction term
