## Supplemental Tables for "Locus coeruleus degeneration is associated with cortical tau deposition and cognitive decline in older adults at familial risk of Alzheimer’s disease"

S. Table 1 | Model statistics for the relationship between LC degeneration and cognitive decline. Each outcome variable was run as a separate model, with all main effects of interest listed. Bolded and italicised main effect titles indicate primary hypotheses. Statistics shown in bold highlight significant (p<0.05) effects.

| Main effects | Outcome variable | t-value | p-value | df |  | Outcome variable | t-value | p-value | df |
| --- | --- | --- | --- | --- | --- | --- | --- | --- | --- |
| Time | RBANS Total | **-6.92** | **0.000** | **1201** |  | Delayed Recall | **-3.60** | **0.000** | **1224** |
| bl-LC |  | 1.14 | 0.255 | 184 |  |  | 0.42 | 0.675 | 188 |
| LC-slope |  | 1.69 | 0.094 | 190 |  |  | 1.77 | 0.078 | 199 |
| Amyloid |  | **-2.94** | **0.004** | **186** |  |  | **-4.45** | **0.000** | **191** |
| ***bl-LC*Time*** |  | **2.31** | **0.021** | **1217** |  |  | 0.37 | 0.714 | 1247 |
| ***LC-slope*Time*** |  | **2.18** | **0.030** | **1216** |  |  | 1.68 | 0.093 | 1246 |
| Amyloid*Time |  | **-5.35** | **0.000** | **1204** |  |  | **-5.86** | **0.000** | **1229** |
| bl-LC*Amyloid |  | 0.40 | 0.691 | 190 |  |  | 0.08 | 0.938 | 199 |
| LC-slope*Amyloid |  | 0.84 | 0.403 | 200 |  |  | 1.83 | 0.068 | 215 |
| ***bl-LC*Amyloid*Time*** |  | 1.03 | 0.304 | 1224 |  |  | 0.31 | 0.760 | 1256 |
| ***LC-slope*Amyloid*Time*** |  | 1.87 | 0.062 | 1236 |  |  | **4.31** | **0.000** | **1272** |
| Time | Attention | 1.09 | 0.275 | 1186 |  | Language | **-2.75** | **0.006** | **1273** |
| bl-LC |  | 0.99 | 0.323 | 179 |  |  | 1.55 | 0.123 | 216 |
| LC-slope |  | 0.72 | 0.472 | 184 |  |  | 0.15 | 0.882 | 237 |
| Amyloid |  | -1.35 | 0.177 | 180 |  |  | -0.87 | 0.385 | 224 |
| ***bl-LC*Time*** |  | **3.54** | **0.000** | **1199** |  |  | 0.73 | 0.466 | 1296 |
| ***LC-slope*Time*** |  | 0.63 | 0.528 | 1198 |  |  | -0.06 | 0.950 | 1294 |
| Amyloid*Time |  | **-3.83** | **0.000** | **1189** |  |  | -0.84 | 0.400 | 1278 |
| bl-LC*Amyloid |  | -0.71 | 0.477 | 184 |  |  | 0.88 | 0.377 | 238 |
| LC-slope*Amyloid |  | 0.21 | 0.836 | 190 |  |  | 0.70 | 0.485 | 268 |
| ***bl-LC*Amyloid*Time*** |  | 1.00 | 0.320 | 1204 |  |  | 1.70 | 0.090 | 1304 |
| ***LC-slope*Amyloid*Time*** |  | -1.47 | 0.142 | 1213 |  |  | 0.65 | 0.518 | 1316 |
| Time | Immediate Recall | -1.12 | 0.265 | 1239 |  | Visuospatial/Constructional | **-11.76** | **0.000** | **1246** |
| bl-LC |  | 1.16 | 0.246 | 190 |  |  | -0.09 | 0.925 | 199 |
| LC-slope |  | **2.57** | **0.011** | **201** |  |  | 0.69 | 0.489 | 214 |
| Amyloid |  | **-2.83** | **0.005** | **193** |  |  | -1.33 | 0.185 | 204 |
| ***bl-LC*Time*** |  | 0.98 | 0.327 | 1255 |  |  | 1.45 | 0.147 | 1271 |
| ***LC-slope*Time*** |  | **2.86** | **0.004** | **1253** |  |  | 1.26 | 0.209 | 1268 |
| Amyloid*Time |  | **-5.23** | **0.000** | **1236** |  |  | -0.65 | 0.518 | 1251 |
| bl-LC*Amyloid |  | 1.12 | 0.266 | 201 |  |  | 1.04 | 0.300 | 214 |
| LC-slope*Amyloid |  | 1.31 | 0.192 | 219 |  |  | -0.19 | 0.850 | 236 |
| ***bl-LC*Amyloid*Time*** |  | 1.70 | 0.090 | 1264 |  |  | -0.83 | 0.408 | 1280 |
| ***LC-slope*Amyloid*Time*** |  | **2.43** | **0.015** | **1280** |  |  | -0.06 | 0.949 | 1295 |

S. Table 2 | Interpretation guide for main effects listed in S. Table 1.

| Main Effect | Interpretation |
| --- | --- |
| Time | Does this cognitive score change over time? |
| bl-LC | Do cognitive scores at time 0 differ according to bl-LC? |
| LC-slope | Do cognitive scores at time 0 differ according to LC-slope? |
| Amyloid | Do cognitive scores at time 0 differ according to global amyloid? |
| bl-LC*Time | Does rate of cognitive change depend of bl-LC? |
| LC-slope*Time | Does rate of cognitive change depend on LC-slope? |
| Amyloid*Time | Does rate of cognitive change depend on global amyloid? |
| bl-LC*Amyloid | Is any dependence of cognition at time 0 on LC-bl stronger in people with more global amyloid? |
| LC-slope*Amyloid | Is any dependence of cognition at time 0 on LC-slope stronger in people with more global amyloid? |
| bl-LC*Amyloid*Time | Is any dependence of the rate of cognitive change on LC-bl stronger in people with more global amyloid? |
| LC-slope*Amyloid*Time | Is any dependence of the rate of cognitive change on LC-slope stronger in people with more global amyloid? |
